# Combining data independent acquisition with spike-in SILAC (DIA-SiS) improves proteome coverage and quantification

**DOI:** 10.1101/2024.05.03.592381

**Authors:** Anna Sophie Welter, Maximilian Gerwien, Robert Kerridge, Keziban Merve Alp, Philipp Mertins, Matthias Selbach

## Abstract

Data Independent Acquisition (DIA) is increasingly preferred over Data Dependent Acquisition (DDA) due to its higher throughput and fewer missing values. Whereas DDA often utilizes stable isotope labeling to improve quantification, DIA mostly relies on label-free approaches. Efforts to integrate DIA with isotope labeling include chemical methods like mTRAQ and dimethyl labeling, which, while effective, complicate sample preparation. Stable isotope labeling by amino acids in cell culture (SILAC) achieves high labeling efficiency through the metabolic incorporation of heavy labels into proteins *in vivo*. However, the need for metabolic incorporation limits the direct use in clinical scenarios. Spike-in SILAC methods utilize an externally generated heavy sample as an internal reference, enabling SILAC-based quantification even for samples that cannot be directly labeled. Here, we combine DIA with spike-in SILAC (DIA-SiS), leveraging the robust quantification of SILAC without the complexities associated with chemical labeling. We developed and rigorously validated DIA-SiS through a mixed-species benchmark to assess its performance in proteome coverage and quantification. We demonstrate that DIA-SiS significantly improves proteome coverage and quantification compared to label-free approaches and reduces the incidence of incorrectly quantified proteins. Additionally, DIA-SiS proves effective in analyzing proteins in low-input formalin-fixed paraffin-embedded (FFPE) tissue sections. DIA-SiS combines the precision of stable isotope-based quantification with the simplicity of label-free sample preparation, facilitating simple, accurate and comprehensive proteome profiling.

## Introduction

In mass spectrometry-based proteomics, Data-Independent Acquisition (DIA) is gaining popularity over Data-Dependent Acquisition (DDA) for its higher throughput and fewer missing values (1–6). Quantification in DIA typically utilizes label-free methods (7–9). However, since stable isotope-based quantification methods offer superior quantification in DDA (10), efforts have been made to extend those techniques to DIA. Chemical stable isotope labeling methods such as mTRAQ and dimethyl labeling have been successfully combined with DIA (11, 12). However, chemical labeling requires optimization to achieve high efficiency, and the additional *in vitro* steps complicate sample preparation and increase variability. *Stable isotope labeling by amino acids in cell culture* (SILAC) achieves high labeling efficiency through metabolic incorporation of heavy labels into proteins *in vivo* (13). This simplifies sample preparation since no extra chemical labeling steps are required. Moreover, combining differentially labeled samples early during sample preparation eliminates any variation that might be result from sample processing. Due to these advantages, SILAC has become a popular method for functional proteomics, especially in combination with DDA (14, 15). As a metabolic labeling method, SILAC furthermore enables quantification of protein synthesis and degradation (16–19). Recent studies successfully combined SILAC with DIA to study protein turnover (20–23).

The primary limitation of SILAC is the need for metabolic incorporation of stable isotope-labeled amino acids, rendering it unsuitable for direct application in clinical scenarios (e.g., human tissues). To address this issue, internal standard or ‘spike-in’ methods have been devised (24, 25). These methods use SILAC to produce a heavy reference sample, which is subsequently added to each unlabeled (light) sample. The ratios of light to heavy proteins are determined in each sample. The uniform heavy reference across all samples serves as an internal standard, enabling relative quantification of light proteins across samples by computing the ratio of these ratios. In this way, spike-in SILAC allows SILAC-based quantification for samples that cannot be directly labeled. A major advantage of spike-in SILAC is its decoupling of labeling from sample preparation. Once the heavy spike-in reference is prepared and added, further processing requires no additional labeling, streamlining the workflow. Hence, spike-in SILAC combines the accuracy of stable isotope-based quantification with the simplicity of label free sample preparation. In combination with DDA, this methodology has been successfully implemented in both pre-clinical and clinical studies (26–30).

We reasoned that combining DIA with spike-in SILAC (DIA-SiS) would be a simple technique for sensitive, accurate and precise quantification on a proteome-wide scale. First, DIA-SiS should provide better quantification than label-free approaches. Second, the addition of the heavy spike-in reference could facilitate detection of low abundant proteins. Third, the method is simple since it does not require additional chemical labeling steps. Here, we developed and rigorously benchmarked DIA-SiS on a mixed species dataset with ground truth relative protein abundances. Our results show that DIA-SiS indeed provides better quantification and improves the coverage, especially of low abundant proteins. In this way, DIA-SiS detects differentially abundant proteins with greater sensitivity and specificity. Finally, we show that DIA-SiS boosts protein detection in low input formalin fixed paraffin embedded (FFPE) tissue sections. In summary, these data show that DIA-SiS significantly improves proteome coverage and quantification compared to label free approaches.

## Material and Methods

### Sample preparation

#### Two-species benchmarking experiment

The *E. coli* strain AT713, which cannot synthesize lysine and arginine, was grown in a medium containing 0.5 % (w/v) D-(+)-glucose (Sigma), 1.3 % (w/v) M9 salts (BD), 1 mM MgSO4 (Merck), 377 µM thiamine (Sigma), 300 µg/mL L-arginine (Sigma) and L-lysine (Sigma), plus 150 µg/mL of the 18 other natural amino acids (Sigma). For heavy or light SILAC labeling, Arg10(^13^C_6_,^15^N_4_) and Lys8(^13^C_6_,^15^N_2_) or Arg0 and Lys0 (Silantes) were employed, respectively. To culture the bacteria, the glycerol stock was streaked onto 1.2 % (w/v) lysogeny broth (LB, Sigma) agar plates and grown overnight. Then, single colonies were selected for overnight pre-culture in the defined SILAC light or heavy media. The pre-cultures were then used to inoculate overnight batch cultures, again using the SILAC media. The colonies that were selected for this were tested for arginine and lysine auxotrophy on SILAC medium agar (1.2 % w/v) plates supplemented with 1) nothing, 2) 300 µg/mL arginine, 3) 300 µg/mL lysine, 4) 300 µg/mL arginine and lysine. Growth was only observed in condition 4.

The bacterial cells were harvested by pelleting (1500 rcf, 4 °C, 10 m), then washed with ice-cold PBS (phosphate buffered saline, pH 7.4, ThermoFisher Scientific). The lysis buffer (1 % SDC (Sigma), 150 mM NaCl (Roth), 100 mM Tris (Roth) pH 8, 1 Roche cOmplete protease inhibitor tablet per 10 mL) was added to the pellet to a target protein concentration of 1 mg/mL, vortexed and incubated for 10 m at 96 °C in a thermal cycler (TC). The mixture was frozen at −72 °C in a EtOH/dry ice slush and thawed at 30 °C, shaking at 1000 rpm in a TC. The freeze-thaw cycle was repeated three times in total. The DNA was digested by incubation with benzonase (Sigma, 25 U per expected 1 mg of protein) for 30 min at room temperature. Finally, the lysate was cleared for 15 m at 10,000 rcf and 4 °C. The protein concentration was determined by a BCA assay (Pierce BCA Kit, Thermo Fisher Scientific). The disulfide bridges in the protein extract were reduced in 10 mM dithiothreitol (DTT, Sigma) for 1 h at 37 °C and shaking at 1000 rpm in a TC. The free thiols were then alkylated by adding chloroacetamide (CAA, Sigma) to a concentration of 20 mM and incubating for 45 m at room temperature in the dark (1000 rpm, TC) and the reaction subsequently quenched by increasing DTT to 50 mM. The sample cleanup and digestion into peptides was performed by protein aggregation capture on magnetic beads, based on (31). Briefly, 15:1 (w/w) magnetic beads (Cytiva Speedbeads 45152105050250 and 65152105050250) to expected protein amount were added to the samples. Then, acetonitrile concentration was increased to 75 %. The beads with the proteins bound to them were then washed three times with 80 % EtOH (Chemsolute) and then digested overnight at 37 °C and shaking at 1000 rpm (TC) with trypsin (Sequencing Grade Modified Trypsin V5113, Promega) and LysC (Lysyl Endopeptidase 129-02541, Wako Chemicals) in a 1:50 protease to substrate ratio in ammonium bicarbonate buffer (Sigma, 50 mM). The next day, the peptides were lyophilized in a vacuum concentrator and resolved in solvent A (3 % (v/v) acetonitrile (Chemsolute) and 0.1 % (v/v) formic acid (Fluka)).

Human HL-60 cells were grown at 37 °C and 5 % CO_2_ in T75 flasks. The cells were cultured at a density of 0.5 - 1.5 million cells per mL and pelleted at 300 rcf for 5 min for passaging. The unlabeled (light) cells were grown in RPMI Medium 1640 (Gibco, Thermo Fisher Scientific) supplemented with 10 % fetal bovine serum (FBS, PAN Biotech) and non-essential amino acids, sodium pyruvate as well as L-alanyl-L-glutamine (all Gibco, ThermoFisher Scientific, according to the manufacturer’s instructions). For heavy SILAC labeling, the cells were grown for 6-8 passages in SILAC RPMI Medium (PAN Biotech, P04-02504) which lacks lysine and arginine. It was supplemented with heavy lysine and arginine (Lys8(^13^C_6_,^15^N_2_) and Arg10(^13^C_6_,^15^N_4_), Silantes), 10 % dialyzed FBS (PAN Biotech), L-alanyl-L-glutamine (Gibco, ThermoFisher Scientific), non-essential amino acids (Gibco, ThermoFisher Scientific) and sodium pyruvate (Gibco, ThermoFisher Scientific). Both labeled and unlabeled cells were harvested by pelleting for 5 min at 500 rcf and washed with ice-cold PBS (pH 7.4, ThermoFisher Scientific).

For lysis, the cells were then resuspended by vortexing in a buffer composed of 1 % (w/v) SDS (Roth), 5 % (v/v) ACN (Chemsolute) in PBS (pH 7.4, ThermoFisher Scientific) to a target concentration of 1 mg/mL. They were incubated for 10 min at 96 °C and subsequently sonicated in a Bioruptor plus ultrasonicator for 10 cycles (30 s on/30 s off, 4 °C) on the high intensity sonication setting. Then, the lysates were cleared by centrifugation (15 m, 10,000 rcf, 4°C) and the protein concentration determined by a BCA assay (Pierce BCA Kit, Thermo Fisher Scientific). The human protein extracts were further processed to peptides in the same manner as described above for bacterial protein extracts.

The peptides were then mixed according to Figure 1A and the injection amounts are indicated in the cartoon. For the low input samples (Figure 3D-G), 13.4 ng light *H. sapiens* peptides with either 13.4 ng light *E. coli* peptides or no light *E. coli* peptides, plus as a spike-in 66.8 ng heavy *H. sapiens* peptides with 6.7 ng heavy *E. coli* peptides were injected.

**Figure 1:**
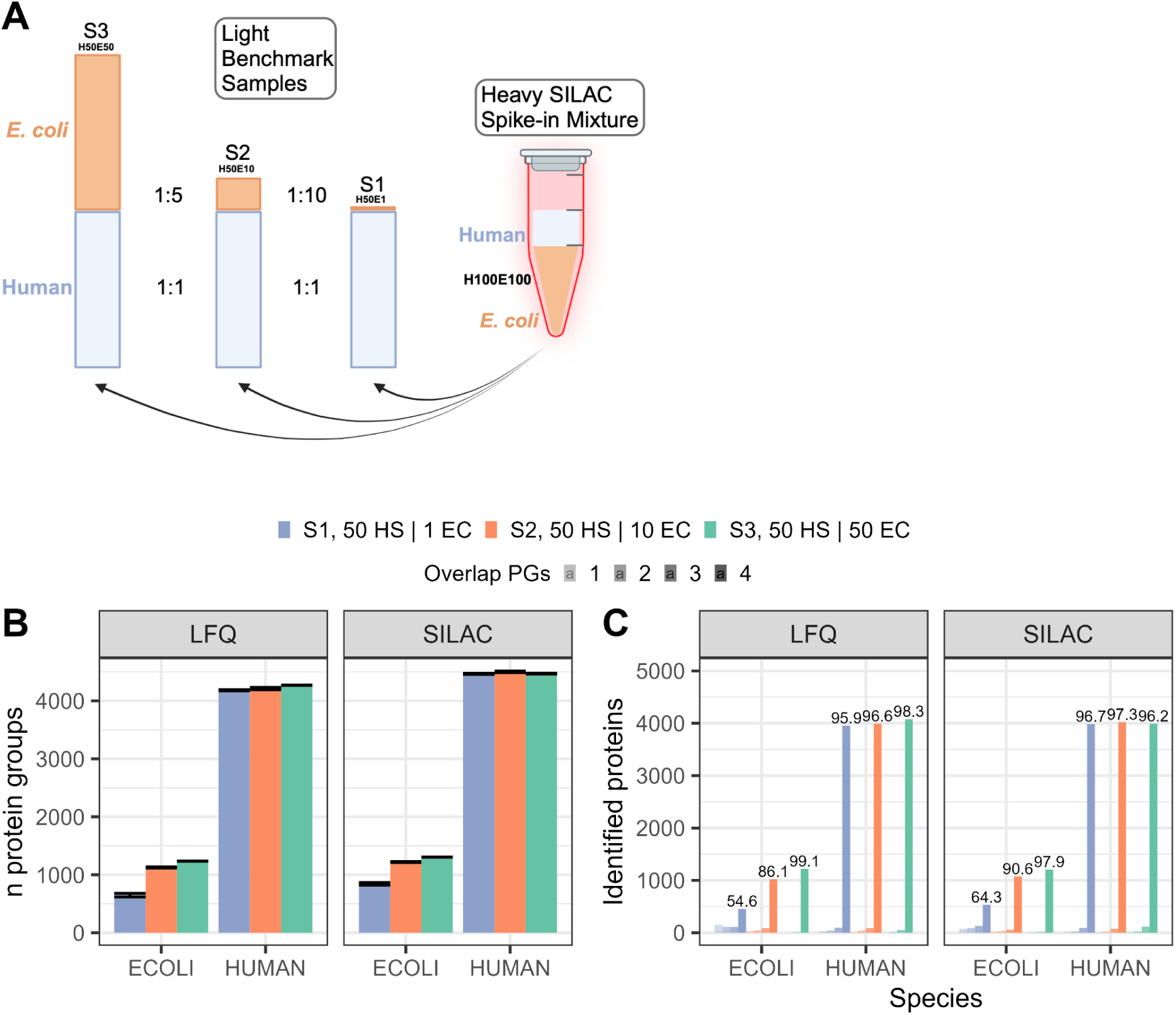
Benchmark experiment: Design and protein coverage. **A)** Experimental design of the mixed species benchmark. **B)** Mean number of identified *E.coli* and human HL-60 proteins over all 4 technical replicates per dilution step sample (standard deviation is indicated by whiskers). The different colors correspond to different *E.coli* dilutions. **C)** Number of identified proteins one, two, three or all four replicates (increasing opacity). Numbers indicate the percentage of human or *E. coli* protein groups detected in all 4 replicates.

**Figure 2:**
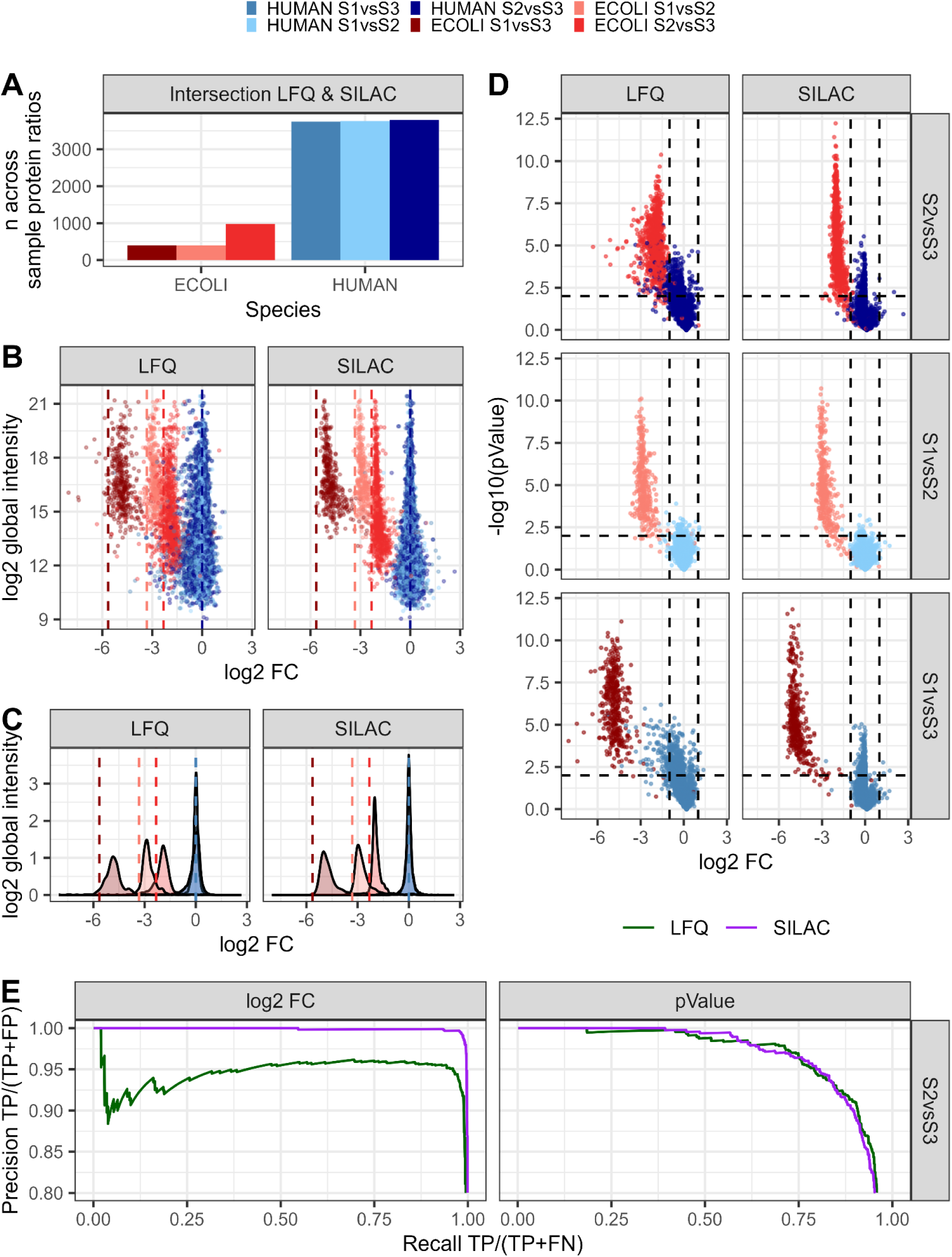
DIA-SiS allows reliable across-sample quantification. Only proteins with both LFQ and SILAC ratios and no missing values across all replicates are shown. **A**) The bars indicate the number of across-sample protein ratios quantified with both LFQ and DIA-SiS. **B)** Both LFQ and DIA-SiS capture the expected across-sample ratios, with a clearer intensity-dependent precision for DIA-SiS. The global protein group abundance is plotted against the mean across-sample protein ratios. Dashed lines indicate expected ratios. **C)** Density plots corresponding to B). **D)** DIA-SiS improves detection of differentially abundant proteins. The volcano plots show the log2FCs on the x axis and the −log10 p-values on the y axis. The dashed lines indicate cutoffs (p-value = 0.01, absolute log2FC = 1), blue: human proteins, red: *E. coli* proteins. **E)** Precision-recall curves based on D) of S2 vs S3 for log2FCs (left) and p-values (right) for LFQ (green) and SILAC (purple).

**Figure 3:**
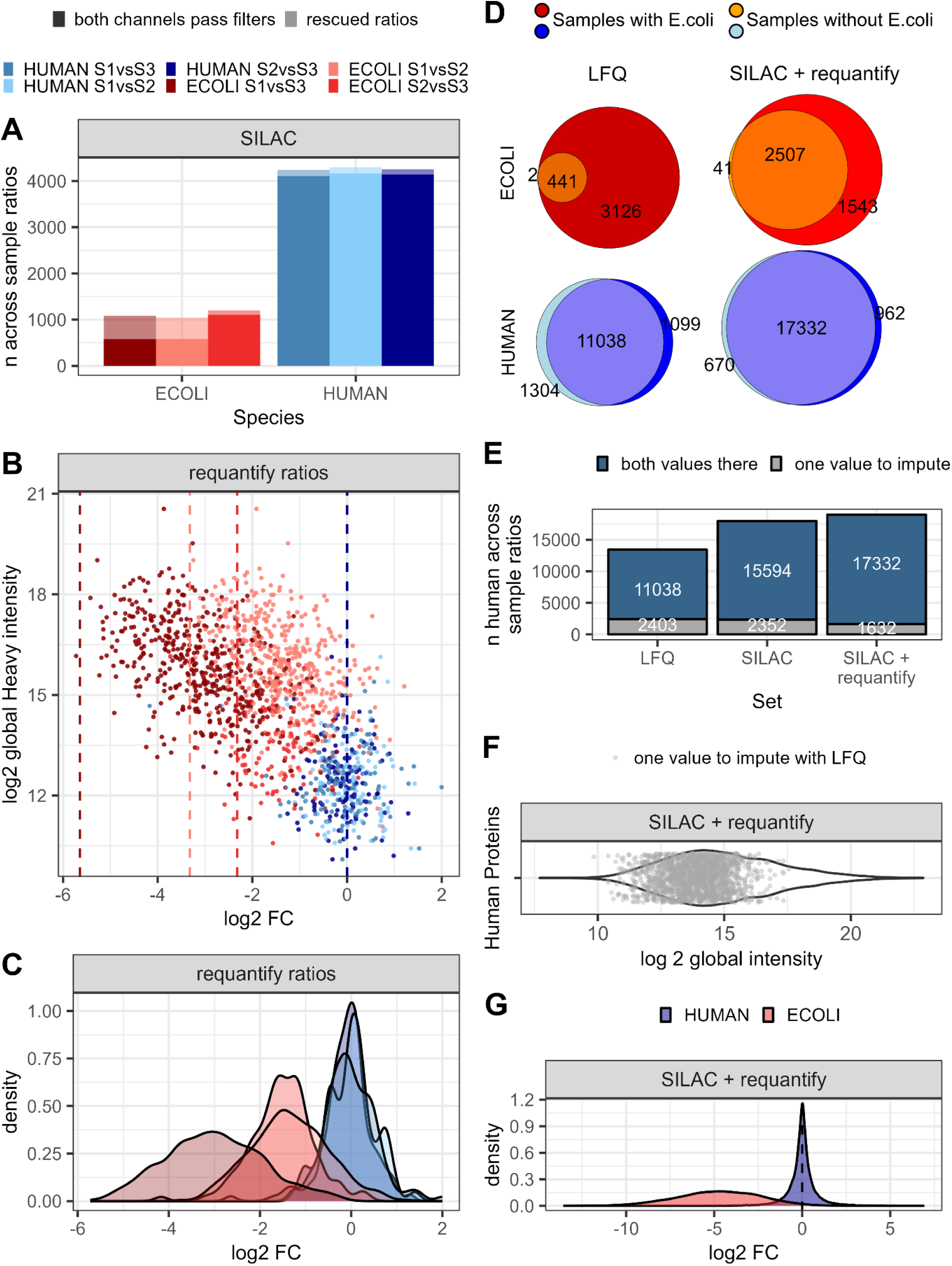
“Requantify” further reduces missing values in DIA-SiS. **A)** The number of proteins quantified across samples can be increased by only requiring confident identification of heavy channel precursors (“requantify” option). Mean number of across sample ratios (without missing values) using DIA-SiS, requiring the light as well as heavy channel to pass filters (higher opacity) and additional ratios obtained when using requantify (lower opacity). **B)** Requantified DIA-SiS ratios still capture the correct trend. The log global abundance is plotted against the mean log2FCs of protein groups (blue: human proteins, red: *E. coli* proteins). **C)** Density plots corresponding to B); **D-G)** Comparison of a sample containing human and *E. coli* proteins to sample w/o *E. coli* shows that DIA-SiS reduces missing values, especially with “requantify” enabled. **D)** Cumulative Venn diagrams of *E.coli* (upper, red/orange) and human (lower, blue) proteins quantified in four technical replicates. **E)** DIA-SiS also reduces missing values for human proteins although they were equally abundant in both samples. Cumulative bar charts for the human proteins illustrated in D) indicating the number of ratios that could be computed between samples. **F)** Rescued human proteins cover a broad abundance range. The cumulative distribution shows the log2 global intensity of those proteins that are missing in LFQ but could be rescued with DIA-SiS and requantify. The violin indicates the intensity of the entirety of proteins found with SiS and requantify. **G)** Across-sample ratios obtained using SILAC + requantify correctly capture the global trend (blue: human, red: *E.coli*).

#### Super-DIA-SiS

For the super-SILAC spike-in mix, six different HPV negative head and neck squamous cell carcinoma cell lines (A-253 (salivary gland), Cal-33 (tongue), FaDu (pharynx), UT-SCC-14 (tongue), SCC-25 (tongue), UPCI-SCC-026 (tongue)) were grown at 37 °C and 5 % CO_2_ in 10 cm and 15 cm (last two passages) dishes. After thawing, cells were kept on DMEM Medium (Gibco, Thermo Fisher Scientific) supplemented with 10 % FBS (PAN Biotech) for two to three passages until they recovered. After recovery, the medium was exchanged to SILAC DMEM Medium (Pan Biotech, P04-02501) supplemented with 10 % dialyzed FBS (Pan Biotech), L-alanyl-L-glutamine (Gibco, ThermoFisher Scientific, according to the manufacturer’s instructions) and heavy lysine and arginine (Lys8 and Arg10, Silantes). In order to receive maximal labeling efficacy, cells were split 1:4 at 70 to 80 % confluency six times over the course of five to six weeks. At each passage, cells were washed with PBS (pH 7.4, ThermoFisher Scientific) and incubated with 2 ml 0.05 % Trypsin EDTA (Gibco, ThermoFisher Scientific) for 10 to 15 min to detach the cells. Trypsin was blocked by adding 2 ml of medium. To avoid light lysine and arginine contamination from Trypsin EDTA, cells were pelleted for 5 min at 300 rcf and resuspended in the new medium.

For harvesting, the dishes were washed three times with ice-cold PBS (pH 7.4, ThermoFisher Scientific) and transferred to microcentrifuge tubes using cell scrapers. The cell suspension was centrifuged at 16,000 rcf for 15 min at 4 °C. Cell pellets were stored in −80 °C freezer until further usage. For the lysis, the pellets were suspended in lysis buffer (50 mM TRIS (Roth) pH 8, 1 % SDS (w/v) (Roth), 150 mM NaCl (Roth), 1 Protease Inhibitor tablet (Roche cOmplete) per 10 ml) and boiled for 5 min at 95 °C. Lysates were ultra sonicated (Bioruptor Plus) for 10 cycles (30 s on / 30 s off) on high intensity sonication setting and centrifuged at 16,000 rcf for 15 min at RT to pellet the cell debris. Protein concentration of each lysate was measured via BCA (Pierce BCA Kit, Thermo Fisher Scientific).

HNSCC FFPE samples were kindly provided by Dr. Ingeborg Tinhofer-Keilholz (Laboratory for Translational Radiooncology, Dept. of Radiooncology and Radiotherapy, Charité Universitätsmedizin Berlin). Deparaffinization was done following a partially modified protocol from the High Pure FFPET DNA Isolation Kit (Roche). Briefly, FFPE slices were bathed in xylene (10 min at RT) and twice in fresh absolute ethanol (10 min at RT) and scraped into a tube. Samples were then centrifuged to pellet the tissues at 16,000 g for 2 min at RT. Pellets were washed by adding 70 % ethanol, five seconds of vortexing and pelleting the tissue again at 16,000 g for 4 min at RT. Ethanol was removed, ultrapure water was added and the solution centrifuged again at 16,000 g for 10 min at RT. The pellets were stored at −80°C until further use. For the lysis a lysis buffer (4 % (w/v) SDS (Roth), 25 mM Tris (Roth), 2.5 mM DTT (Sigma)) was added and the samples were incubated at 900 rpm for 2 hours at 95 °C. The samples were then sonicated with a Covaris LE220Rsc at 250 W for 5 min. The samples were subsequently transferred back into microcentrifuge tubes and incubated at 900 rpm for 90 min at 95 °C. Pellet and suspension were then separated at 12700 rpm for 30 min. The protein concentration was measured via BCA (Pierce BCA Kit, Thermo Fisher Scientific).

The FFPEs were either processed alone (300 ng and 50 ng samples) or together with the spike-in (50 ng + spike-in). To this end, equal amounts of the six different heavy labeled cell lysates were mixed to generate a super-SILAC spike-in mixture. This mixture was added to the FFPE extracts in such a way that the protein amount of the spike-in was five times the amount of FFPE proteins. The samples were processed (reduction, alkylation, SP3) in three batches in a semi-automated manner in a 96 well plate using an Opentrons OT2 (Opentrons Labworks Inc.). Briefly, samples were reduced by incubating with 10 mM DTT (Sigma) for 30 min at RT shaking and alkylated with 20 mM Iodoacetamide (Sigma) for 30 min at RT shaking. Alkylation was quenched by adding DTT (Sigma) to a final concentration of 50 mM and incubation for 15 min at RT shaking. The protein cleanup and digestion steps followed the same protocol as described for the benchmarks besides using a 20:1 bead to protein ratio. After digestion, peptides were further cleaned utilizing a G5563A Bravo platform (Agilent). Briefly, the syringes of the robot’s pipetting head were washed with a washing buffer (50% ACN (Chemsolute) and 0.1% TFA (Sigma)) to flush potential contaminants. The same buffer was used to condition the resin of a AssayMAP C18 cartridge rack. A solution of 0.1 % TFA (Sigma) was run through the cartridge to equilibrate them. Afterwards, the samples were loaded onto the resin. Syringes and cartridge were washed with the washing buffer before eluting the samples in a fresh 96-well plate using the washing buffer. Afterwards, the peptides were lyophilized in a vacuum concentrator and resolved in solvent A (3 % (v/v) acetonitrile (Chemsolute) and 0.1 % (v/v) formic acid (Fluka)).

### Labeling efficiency

For the labeling checks of the heavy labeled cells, roughly 500 ng peptides were injected in a Orbitrap Exploris 480 (Thermo Fisher Scientific) coupled to a Vanquish Neo UHPLC-System (for Super-DIA-SiS; Thermo Fisher Scientific) or an EASY-nLC 1200 (for benchmarks; Thermo Fisher Scientific) running in nanoflow mode. The attached column was an in-house packed 20 cm 1.9 µm column. Samples were measured in DDA mode utilizing a 44 min (Super-DIA-SiS) or a 110 min gradient (benchmarks). Raw files were analyzed with MaxQuant 1.6.7.0. The evidence files were filtered for peptides containing one arginine or lysine and sorted after their intensity. Some of the highest intensity precursors with low PEP were visualized using Thermo Xcalibur 4.6.67.17 Qual Browser (Thermo Fisher Scientific) and Inkscape 1.1.

### Liquid Chromatography and Mass spectrometry

All DIA measurements were performed on a timsTOF Pro 2 (Bruker Daltonics) attached to a EASY-nLC 1200 (Thermo Fisher Scientific) in DIA-PASEF mode utilizing an in-house packed column (20 cm long, 75 μm diameter, 1.9-μm ReproSil-Pur C18-AQ silica beads, Dr Maisch) heated to 50 °C connected to a nano-electrospray ion source (CaptiveSpray; Bruker Daltonics) with a spray voltage of 1600 V. The peptides were separated using a ∼ 30 min active gradient at a constant flow rate of 250 nL/min from 0 % solvent B to 90 % solvent B (0 min, 2 %; 1 min, 7 %; 20 min 20 %; 29 min 30 %; 32 min 60 %; 33 min 90 %). Solvent A consisted of 3 % acetonitrile (LC-MS grade, Chemsolute) and 0.1 % formic acid (Fluka) in H_2_O (LC-MS grade, Chemsolute) and Solvent B of 90 % acetonitrile (LC-MS grade, Chemsolute) and 0,1 % formic acid (Fluka) in H_2_O (LC-MS grade, Chemsolute). The acquisition was operated in DIA-PASEF (32) mode with the mass range set to 400 - 1200 m/z, mobility range set to 0.60 - 1.60 V*s/cm^2^, a ramp time of 100 ms and an accumulation time of 100 ms with an estimated cycle time of 1.80 s and a collision energy of 10 eV (’DIA-PASEF long gradient’, as provided by Bruker Daltonics). To check the SILAC labeling efficiency, samples were measured in DDA mode on an Orbitrap Exploris 480 (Thermo Fisher Scientific) coupled to a Vanquish Neo UHPLC-System (Super-DIA-SiS; Thermo Fisher Scientific) or an EASY-nLC 1200 (benchmarks; Thermo Fisher Scientific) with an electrospray ionization source (Thermo Fisher Scientific) using same in-house packed columns as described above for reverse phase separation. Over a ∼ 30 min (Super-DIA-SiS) or ∼ 90 min (benchmarks) active gradient and a constant flow rate of 250 nL/min peptides were eluted (30 min active gradient: 0 min, 2 %; 1 min, 7 %; 20 min 20 %; 29 min 30 %; 32 min 60 %; 33 min 90 %; 90 min active gradient: 0 min, 2 %; 1 min, 4 %; 68 min 20 %; 88 min 30 %; 98 min 60 %; 99 min 90 %). The elution solvent A and B composition was the same as for the DIA measurements. Standard MS settings were used, which are briefly: 1) 30 min active gradient Top20: MS1: 60k Orbitrap resolution, 300 % normalized automatic gain control (AGC) target, 10 ms maximum injection time and 350-1600 m/z scan range. For MS2, isolation width was 1.3 Da, 28 % normalized HCD collision energy, Orbitrap resolution 15k, 100 % AGC target with 22 ms maximum injection time and 20 s dynamic exclusion with 10 ppm mass tolerance; 2) 90 min active gradient Top20: MS1: 60k Orbitrap resolution, 300 % normalized automatic gain control (AGC) target, 10 ms maximum injection time and 350-1600 m/z scan range. For MS2, isolation width was 1.3 Da, 28 % normalized HCD collision energy, Orbitrap resolution 15k, 100 % AGC target with 22 ms maximum injection time and 20 s dynamic exclusion with 10 ppm mass tolerance.

### Data Analysis

Raw files were processed with DIA-NN v 1.8.1. In order to increase the analysis speed, we generated specific refined libraries for the benchmark and the Super-DIA-SiS experiments. For the mixed species benchmarks, FASTA files for *E. coli* (UP000000625, accessed 2023/01/25), *H. sapiens* (UP000005640, accessed 2023/01/25) and contaminants (Universal Contaminant Protein FASTA from REF-Protein contaminants matter) were concatenated and used for the generation of a predicted library using the following DIA-NN settings: --fasta-search --min-fr-mz 200 --max-fr-mz 1800 --met-excision --cut K*,R* --missed-cleavages 1 --min-pep-len 7 --max-pep-len 30 --min-pr-mz 300 --max-pr-mz 1800 --min-pr-charge 1 --max-pr-charge 4 --unimod4 --reanalyse --relaxed-prot-inf --smart-profiling --peak-center --no-ifs-removal. The resulting predicted library was then used as the basis for library refinement. For the benchmarks, the refinement was based on quadruplicates of the label-free samples (using the lowest dilution, i.e., the samples with the largest amount of *E. coli* present). For library refinement, DIA-NN settings were left at default besides changing the library generation setting to ‘IDs, RT & IM profiling’.

For the generation of the predicted library for our Super-DIA-SiS approach, the same *H. sapiens* and contaminants FASTA files as used in the two-species benchmark experiment were combined and used to predict the library. This library was refined using the first technical replicate per FFPE sample of the high input label free measurements (300 ng). All samples then have been analyzed using the corresponding refined libraries. For the analysis of the label-free samples, the default DIA-NN settings have been used plus the additional option “--no-norm” for the benchmarks. For the SILAC analysis in DIA-NN, the following settings have been changed / added: switch MBR off, --fixed-mod SILAC,0.0,KR,label, --lib-fixed-mod SILAC, --channels SILAC,L,KR,0:0;SILAC,H,KR,8.014199:10.008269, --peak-translation, --original-mods, --no-norm (only for benchmarks), --no-maxlf, --peak-center.

Data analysis and most of the figures were done in R 4.2.2 in RStudio 2022.12.0 Build 353, additionally using packages *tidyverse* 2.0.0 (*33*), *RColorBrewer* 1.1.3 (34)*, ComplexUpset* 1.3.3 (http://doi.org/10.5281/zenodo.3700590, (35)*, pROC* (*36*), *eulerr* 7.0.1 (37) and *ggpubr* 0.6.0 (38). Experimental designs have been generated with Biorender and figures have been polished using Affinity Designer 2 2.3.0. R code for the manuscript as well as the general DIA-SiS pipeline and Python code for the DIA-SiS pipeline is available on github (DOIs will be provided upon acceptance).

The report.tsv files from the DIA-NN output were used for further data analysis. Label-free (LFQ) reports were filtered for contaminants, Precursor.Charge > 1, Lib.PG.Q.Value < 0.01 and Lib.Q.Value < 0.01. The column PG.MaxLFQ was used as quantitative output for each protein group. SILAC reports were filtered for contaminants and Precursor.Charge > 1. Additionally, Global.PG.Q.Value < 0.01 and Channel.Q.Value < 0.03 were used as filters for a) both, the light and heavy channel (basic filtering) and b) only the heavy channel (requantify). For requantify, if only the heavy channel passed q value filtering, the light precursor was allowed to pass as well.

Precursor SILAC ratios were calculated by dividing the light ‘Ms1.Translated’ or ‘Precursor.Translated’ precursor intensities by the corresponding heavy partner. After log10 transformation, protein level L/H ratios were calculated by taking the median of all ‘Ms1.Translated’ and ‘Precursor.Translated’ L/H ratios for all precursors of a given protein. Light protein abundances have been calculated by multiplying (i.e., adding, in log space) the protein L/H ratio with the global median heavy intensity of that protein. The global median heavy intensity was calculated by 1) summing up all precursor intensities (Ms1.Translated and Precursor.Translated) for each protein and 2) taking the median log10 of these summed intensities across all samples. In order to normalize the Super-DIA-SiS data, protein L/H ratios in one run were shifted by the median of all L/H protein ratios of that run before calculating the abundances. The benchmark data was not normalized at this point.

Across-sample-ratios were calculated by dividing protein L/H ratios (spike-in data) or LFQ (label-free data) values of two samples. In order to assess the accuracy in the benchmarking experiments, all across-sample ratios were median normalized based on the human 1:1 ratios. To do so, we calculated the difference between the theoretical (i.e., 1:1) and observed median human log2FC and shifted all *E. coli* and human proteins by this difference. For the across-sample comparisons of log2FCs, proteins with missing values in any replicate were ignored.

For the differential abundance analysis in Fig. 2 and 4, we performed two-sided t-tests without multiple testing correction. For Fig. 2 D proteins with a p-value < 0.01 and an absolute log2FC > 1 was considered significantly differentially abundant while for Fig. 4 B only the p-value cutoff was used for the 300 ng samples. To generate the Precision-Recall curves (Fig. 2 E), only proteins with negative fold-changes were used. In order to enable a fair comparison between label-free FFPE data and SiS FFPE data, the proteins were reduced to proteins found in any of the spike-in samples. This reduces the proteins under consideration by ca. 7.5-8.2 % (from 5500-6000 with 300 ng LFQ) and by ca. 6.3-10 % (from 3000-4800 with 50 ng LFQ).

**Figure 4:**
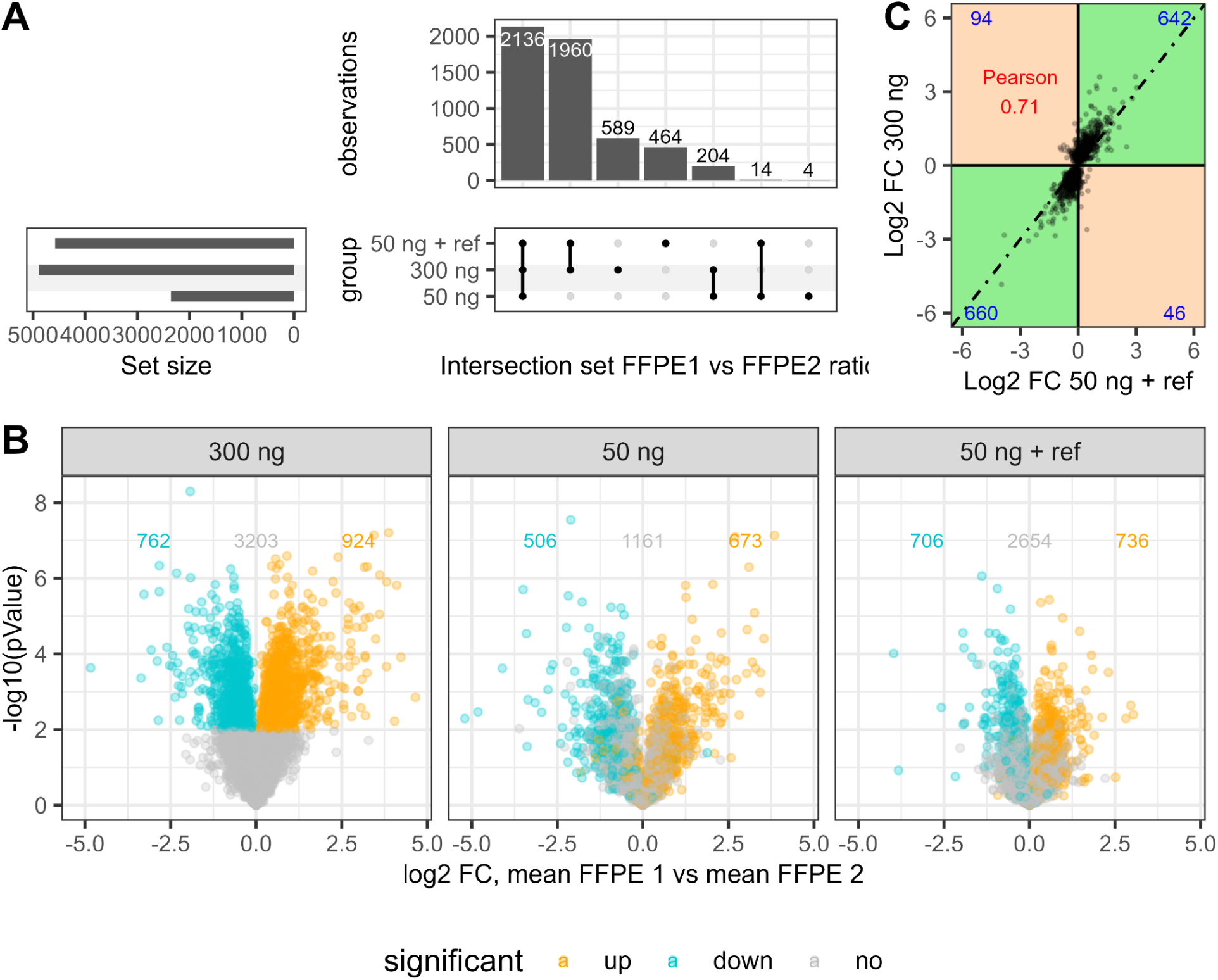
Application of DIA-SiS to formalin-fixed paraffin-embedded (FFPE) head and neck squamous cell carcinoma (HNSCC) samples. Two FFPE tissue samples (FFPE1 and FFPE2) were compared using different input amounts: 300 ng, 50 ng and 50 ng + 250 ng of the SuperSILAC spike-in reference (50 ng + ref). **A)** Number of across sample ratios obtained. Reducing the input from 300 to 50 ng reduces the number of proteins that can be quantified. Adding the spike-in recovers most of the proteins lost in the low input sample. **B)** Volcano plots for differentially abundant proteins in the FFPE1 vs FFPE2 sample for the different input amounts. Significantly differentially abundant proteins (turquoise and orange) were defined based on the 300 ng input (p-value <= 0.01) and colored accordingly in the other samples. The number of proteins in each subset is indicated. **C)** The correlation of the log2FCs of the differentially abundant protein between the 300 ng input and the 50 ng + ref input is high. The number of proteins in the quadrants is indicated.

## Results and Discussion

### DIA-spike-in SILAC (DIA-SiS) improves protein coverage, especially for low abundant proteins

To establish DIA-spike-in SILAC (DIA-SiS), we first generated samples for rigorous benchmarking (Fig. 1 A). A popular method to generate such samples is to combine proteomes of different species in defined ratios (8, 9, 11). Those across-sample ratios constitute a ground truth and can therefore be used to assess the accuracy of a quantitative proteomics method. To this end, we mixed protein extracts from human HL-60 cells and *E. coli*, keeping the amounts of human proteins constant while diluting *E. coli* proteins down to a ratio of 1:50 (Fig. 1 A). To obtain heavy SILAC spike-in samples that contain both heavy human and *E. coli* proteins, we labeled both human HL-60 and *E. coli* cells with heavy stable isotope-encoded lysine and arginine (Lys-8 and Arg-10). Since wild-type *E. coli* is prototrophic for all amino acids, we used the strain AT713 that is unable to synthesize lysine and arginine and has previously been used for SILAC labeling (39). We observed high labeling efficiencies for both human and *E. coli* and added both samples to obtain mixed-species samples with a heavy spike-in reference for either species (Fig. S1 A). Those were then benchmarked against mixed-species samples with the same light proteome composition, but which lack the heavy spike-in. All samples were analyzed in quadruplicates on a Bruker timsTOF Pro2 instrument using 44 min gradients in DIA-PASEF mode (32). We injected the same amounts with respect to the light content of the samples for both the label-free (light only) and SiS (light + heavy spike-in) analyses for a direct comparison of both approaches.

We analyzed the data using DIA-NN, an open-access software that has been widely adopted for DIA proteomics (6, 9). For label free quantification (LFQ), we relied on the MaxLFQ algorithm as implemented in DIA-NN (8). To enable SiS-based quantification we developed a pipeline that takes the DIA-NN output and uses the heavy spike-in as an internal reference for across-sample quantification. Briefly, we first obtain light to heavy (L/H) ratios for every protein in each sample from the median of its corresponding L/H precursor ratios. Next, we derive a global heavy intensity for each protein by taking the median of the summed heavy precursor intensities per protein across samples. Finally, for every protein, we multiply this global heavy intensity with the sample-specific L/H ratios to obtain sample-specific protein intensities. We provide the analysis pipeline in R and python on github (DOI will be made available upon acceptance).

We compared the number of protein groups (PG) identified with DIA-LFQ and DIA-SiS (Fig. 1B). For human proteins the numbers were similar with a mild increase of about 10 % in DIA-SiS. For *E. coli* proteins, on the other hand, we observed a marked increase with SiS, especially for the sample with the smallest amounts of *E. coli* proteins contained. Here, SiS identified about 30 % more PGs than LFQ on average (ca. 900 vs 700). This is expected since the “peak translation” feature of DIA-NN can facilitate the detection of lower abundant light precursors by the presence of more abundant heavy precursors (11). We also investigated the data completeness across replicates. We therefore asked how often a given protein group was identified across the quadruplicate measurements (Fig. 1 C). The majority (> 90 %) of human proteins were consistently detected in all four replicates with either approach. For lower *E. coli* amounts, SiS increases the data completeness. We conclude that LFQ and SiS provide overall similar proteome coverage, with advantages of SiS for low abundant proteins.

### DIA-SiS yields more reliable protein quantification

Next, we compared the quantification of SiS and LFQ. To this end, we computed mean across-sample ratios for each protein across the four technical replicates per sample. For a fair comparison, we first looked at proteins identified in all four replicates in both LFQ and SiS (Fig. 2 A). For this subset, we plotted global protein intensities against the observed log2FCs. Reassuringly, both LFQ and SiS yielded protein quantities that are overall consistent with the expected ratios (Fig. 2 B and C). Also, the variance of the ratios was lower for more abundant proteins with both approaches. However, a closer comparison of LFQ and SiS revealed an overall higher precision of SiS (that is, reduced spread of log2 ratios).

To investigate the practical implications of the differences in the quantitative performance of LFQ and SiS, we used Student’s t-test to identify proteins that are significantly changing across replicates and present the data as volcano plots (Fig. 2 D). While again, both LFQ and SiS were able to distinguish human (not changing) and *E. coli* (changing) proteins based on their fold changes, SiS yielded a better separation of the two protein populations. For example, the small number of human proteins that appeared to be differentially abundant between samples in the SiS analysis typically had large (that is, non-significant) p-values. In contrast, the LFQ analysis resulted in a larger number of human proteins that appeared as differentially abundant, some of them with disturbingly small p-values. At a p-value cut-off of 0.01 and a log2FC cut-off of −1 (dashed lines) in the S1 vs. S3 comparison, LFQ misclassifies 109 human proteins as significantly differentially abundant, while SiS only misclassifies a single protein. These observations are further supported by precision recall curves (Fig. 2 E). Strikingly, SiS leads to substantially fewer erroneous differential abundance as well as p-value calls, improving the specificity. Thus, for the subset of proteins covered by both methods, we conclude that SiS improves quantification, which enables substantially better identification of differentially abundant proteins.

In addition to the subset of proteins quantified by both LFQ and SiS, we also looked at proteins exclusively covered in the SiS analysis (140-190 *E. coli* and 350-400 human proteins, Fig. S2 A). As expected, we found that the precision of these ratios was worse (Fig. S2 B, C) and the number of human proteins erroneously classified as differentially abundant was higher (Fig. S2 D) in this subset than for the intersection. However, the vast majority of SILAC exclusive proteins were still correctly classified. We conclude that DIA-SiS markedly improves the reliability of protein quantification.

### Heavy spike-in based re-quantification further improves the sensitivity of DIA-SiS

The findings presented above indicate that DIA-SiS improves protein detection, especially for the lower abundant *E. coli* proteins. This observation is expected due to the “peak translation” feature of DIA-NN (11). However, we reasoned that this does not yet take full advantage of the spike-in’s potential for pinpointing low-abundance peptides: Since light and heavy precursors have identical ion mobility and very similar LC retention times, detecting the heavy reference implies comprehensive coverage in the LC-MS run where the light precursor is anticipated. Therefore, confidently detecting the heavy spike-in reduces the chance that the absence of a corresponding light precursor is caused by technical issues. Based on this idea and inspired by a similar feature in MaxQuant (40), we implemented a “requantify” functionality in our pipeline. If this option is enabled, only the heavy reference is required to pass our filtering criteria. Any corresponding light precursor signal is then accepted by default, and its intensity values are used for quantification. While the low intensity values rescued in this way are expected to be noisy, we reasoned that they might still allow us to capture the right tendency. Enabling the “requantify” feature nearly doubles the quantifiable number of *E. coli* proteins in the S1 vs S3 comparison (Fig. 3 A). Although these requantified ratios exhibit lower precision and accuracy, they correctly capture the expected trends (Fig. 3 B,C). For instance, the majority of *E. coli* proteins display negative log2 ratios, with the higher dilution (1:50) distinctly separated from the lower dilutions. Hence, integrating this “requantify” algorithm expands the coverage of DIA. Even though the additional data are of lower quality, they constitute valuable additional information that would otherwise be discarded.

Missing values pose a significant challenge in quantitative proteomics (41–43). How to interpret missing values depends on whether a protein escapes detection due to random or technical factors (“missing at random”) or due to its inherently low abundance in a sample (“missing not at random”). In other words, the key question is whether identifying a protein in some samples but not in others should be interpreted as evidence for its lower abundance in the latter. With DDA data, missing LFQ values are typically interpreted as “missing not at random” and therefore imputed. A widely used approach is to use simulated values forming a distribution around the detection limit of actually measured intensities (8). Although DIA offers improved data completeness compared to DDA, the issue of missing values persists.

To evaluate the performance of SiS and LFQ in handling missing values, we conducted an additional benchmark study where one sample contained *E. coli* while the other did not. SiS (with requantify enabled) consistently identified more human and *E. coli* proteins than LFQ (Fig. 3 D). As expected, the differences were especially pronounced for *E. coli* proteins, with SiS identifying over five times more across sample ratios (2,507 vs 441) across all four replicates. Importantly, missing values also impacted human proteins (18 % missing values with LFQ), despite their identical abundance in both samples. This issue is mitigated using SILAC (13.2 % missing values) and further improved with SILAC + requantify (8.6 % missing values; Fig. 3 E). Interestingly, comparing the human proteins missing in LFQ but covered by SiS + requantify to the overall protein intensity distribution reveals that not all of the proteins gained by SiS + requantify are actually of low abundance (Fig. 3 F). We note that imputing these missing values using a distribution around the detection limit of measured intensities would lead to the erroneous conclusion that they are down-regulated in one sample. With SiS and requantify, we significantly reduce the number of missing values and thereby avoid the need for imputation for most proteins. Importantly, across sample ratios for the vast majority of *E. coli* proteins (completely missing in one sample, Fig. 3 G) as well as Human proteins (not changing, independent of the benchmark, Fig. 3 B, C, G) showed the correct trend. Therefore, SiS substantially alleviates the missing value problem.

### Applying DIA-SiS to low input tumor samples

Encouraged by these findings, we aimed to evaluate DIA-SiS in a real-world scenario. Our rationale was to capitalize on the enhanced coverage achieved by DIA-SiS to improve identification of proteins in low-input samples. For this purpose, we investigated formalin-fixed paraffin-embedded (FFPE) samples from head and neck squamous cell carcinomas (HNSCC).

Following the Super-SILAC concept (44), we employed SILAC to label six HNSCC cell lines (Fig. S1 B) and combined their lysates to create a mixed heavy spike-in reference. Unlike the benchmark samples involving two species discussed previously, no ground truth data are available for the FFPE samples. Consequently, we depended on the results of a LFQ analysis performed with standard input amounts and evaluated their recovery in low-input analyses both with and without the heavy spike-in. In this regard, we analyzed injections of 300 ng (standard input), 50 ng (low input), and 50 ng + spike-in (low input + SiS) for two HNSCC FFPE samples. All samples were measured in triplicates and examined using LFQ (300 ng and 50 ng) or our DIA-SiS pipeline with requantification enabled (50 ng + spike-in).

As anticipated, reducing the input from 300 ng to 50 ng significantly decreased the number of across sample ratios between two FFPE samples that could be quantified (Fig. 4 A). Adding the heavy spike-in to the low input sample largely recovered the missing protein ratios, increasing the coverage to approximately 4,500. About 90 % of these ratios overlap with the ratios obtained with standard input samples. In order to verify the relevance of the additional IDs, we performed a differential abundance analysis on the 300 ng samples, labeling all proteins with a p-value < 0.01 as up- or down-regulated, depending on their fold change. 70 % of these significantly differentially abundant proteins are also identified in the 50 ng samples, while adding the spike-in increased the coverage to 86 % (Fig. 4 B). Reassuringly, the correlation of log2FCs in the standard input LFQ sample and the low input SiS sample was high for these differentially abundant proteins (R = 0.71, Fig. 4 C). We conclude that DIA-SiS enhances the proteomic coverage of low-input human tissue samples, thereby presenting a viable option for translational investigations.

## Conclusions

Although stable isotope labeling typically offers superior quantitative performance, most DIA studies to date have utilized label-free quantification (LFQ) due to its simplicity and broad applicability. In contrast, chemical stable isotope labeling techniques such as mTRAQ and dimethyl labeling require additional steps, and metabolic labeling via SILAC is limited to cell culture studies. Here, we demonstrate that integrating spike-in SILAC with DIA combines the superior quantification of stable isotope labeling with the simplicity of label-free sample preparation, enabling straightforward and precise proteome profiling with improved coverage, especially for low-abundance proteins.

One limitation of SiS is that adding the heavy reference reduces the amount of light samples that can be injected into the mass spectrometer. This makes SiS especially attractive in low input scenarios, where the limited capacity is not of concern and the gains are particularly pronounced. For example, DIA-SiS is an attractive method for single cell proteomics (45). In such low-input conditions, the simple sample preparation process becomes an especially advantageous feature (11, 12).

While the ease of SILAC makes it the preferred method for cell culture experiments in DIA, obtaining a suitable heavy reference for human tissue samples can be more challenging. Looking ahead, the broader availability of synthetic proteomes may ease this process (46). Moreover, a further advantage of SILAC is its capability to study proteome turnover. This is an area where the reliable quantification of low-abundant proteins can be instrumental. While we did not explore this topic here, it can be expected to benefit substantially from increased coverage and quantitative performance that DIA-SiS can offer.

## Acknowledgements

We thank Inge Tinhofer-Keilholz (Charité) for generously providing the FFPE samples and Vanessa Deborah Sachse (Charité), Christin Beier, and Mohamed Haji (both MDC) for their assistance with sample processing. We thank Atakan Aydin for cell culture support and Florian Mutschler and Henrik Zauber (both MDC) for insightful discussions. Special thanks to Vadim Demichev (Charité) for guidance on setting up DIA-NN and interpreting its output. This research was funded by the German Ministry of Education and Research (BMBF) under the MSTARS program (grant 16LW0240 to M.S.).

## Author contributions

ASW: Formal analysis, Investigation, Methodology, Software, Visualization, Writing - Review & Editing

MG: Investigation, Methodology, Visualization, Writing - Review & Editing

RK: Software

KMA: Investigation

PM: Funding acquisition, Supervision

MS: Conceptualization, Funding acquisition, Project administration, Supervision, Writing - Original Draft, Writing - Review & Editing

## Competing financial interest statement

The authors declare no competing financial interest.

## Supplement

**Figure S1:**
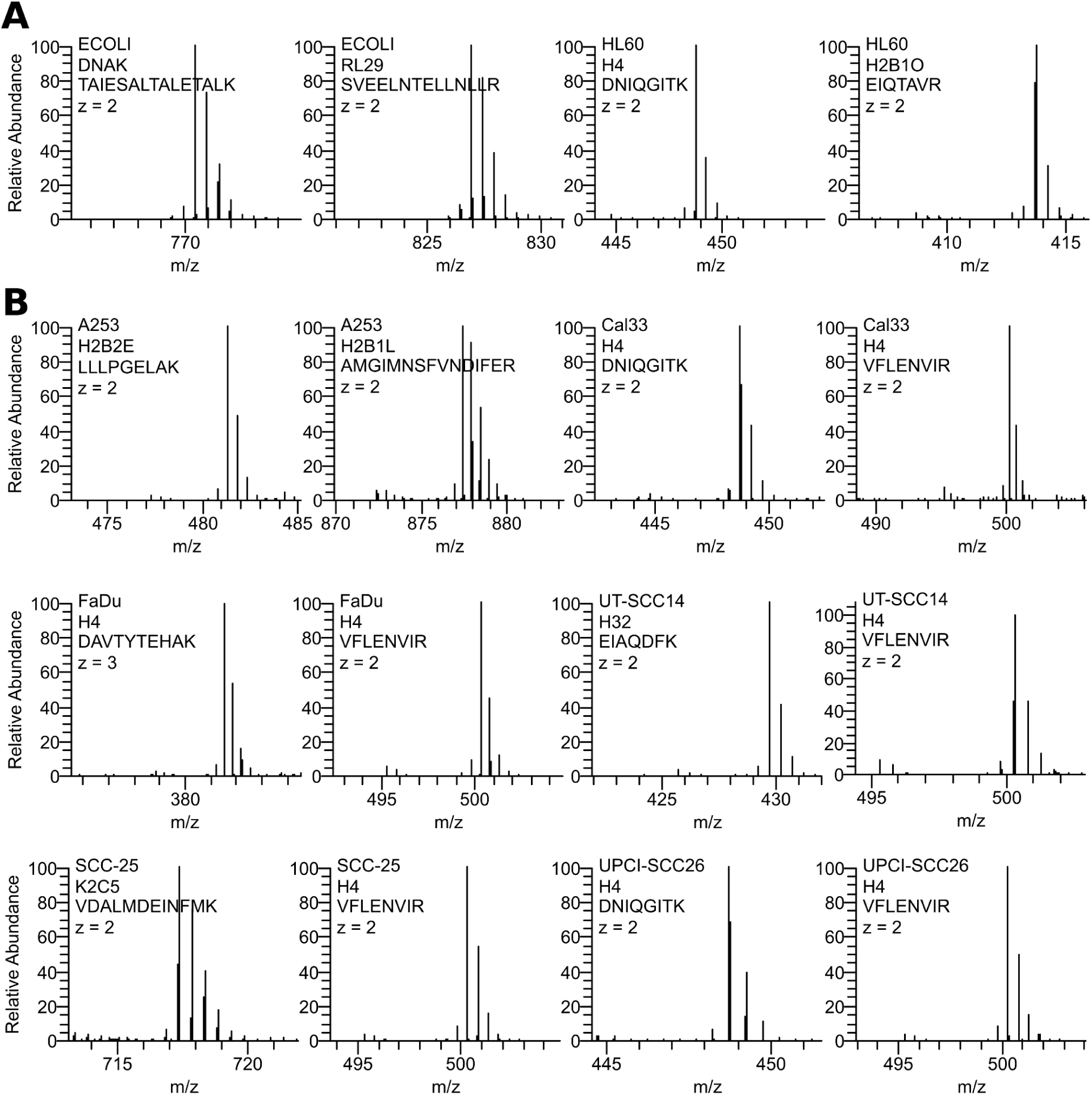
Labeling efficiency of the SILAC spike-ins. Plots show the heavy (100 % relative abundance) and the corresponding light isotopic clusters for one high intensity arginine and lysine containing precursor of all heavy labeled cells used in **A)** the benchmarking experiments and **B)** the Super-DIA-SiS approach.

**Figure S2:**
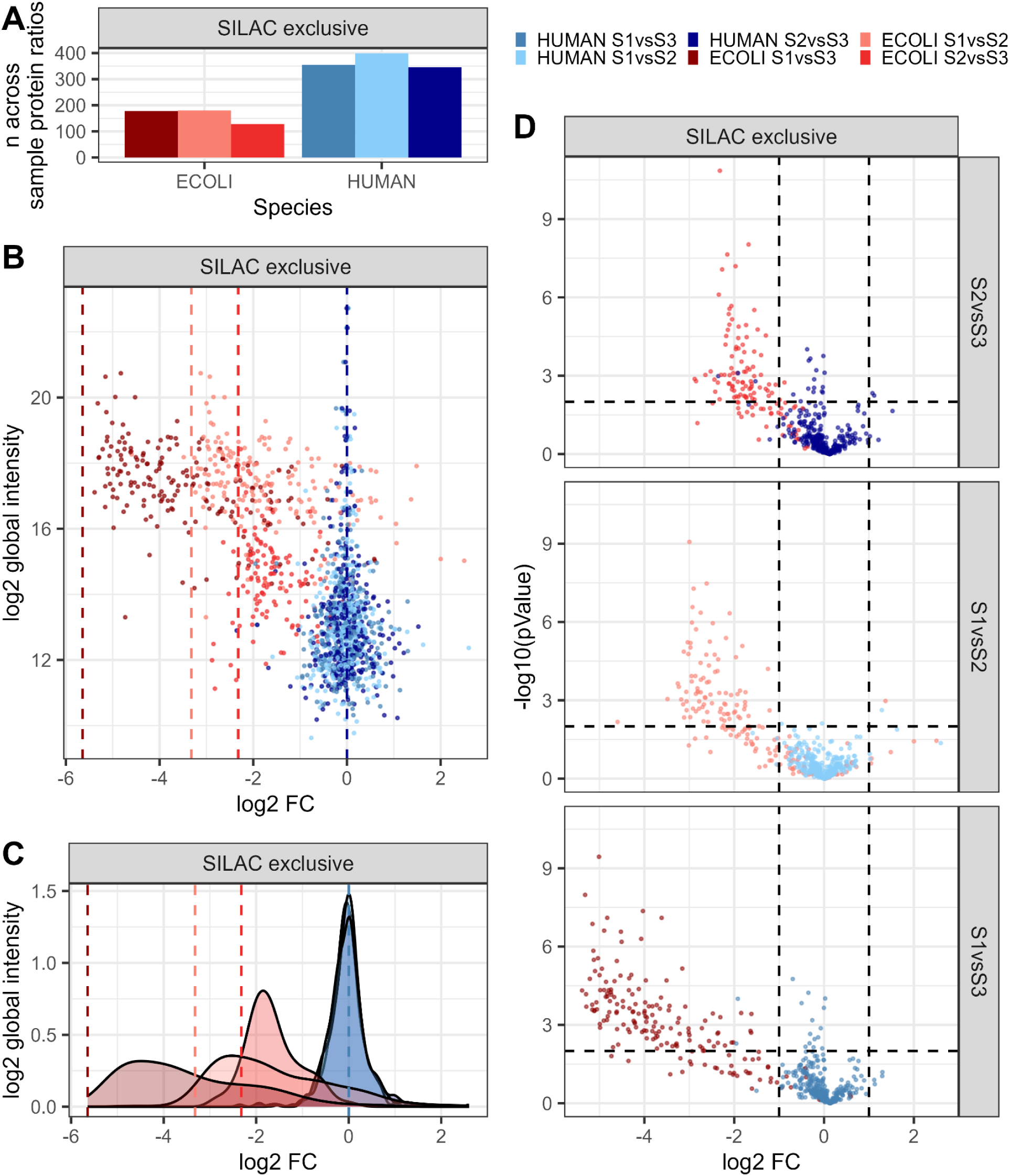
Quantification performance of SILAC-exclusive protein groups. Only proteins with SILAC-exclusive ratios, no missing values across all replicates are shown. **A**) Set size, number of Human (blue) and *E.coli* (red) across-sample protein ratios; **B)** Mean across-sample protein ratios (y-axis) versus global protein group abundance (x-axis). Dashed lines indicate p value cutoffs of > 0.01 and a log2FC of > 1 / < −1, human proteins in blue, *E.coli* in red. **C)** Density of mean across-sample protein ratios (x-axis) corresponding to plots in B); **D)** −log10 p values (y-axis) versus mean across sample ratios (x-axis) for SILAC (right) and three different target ratios (S2vsS3 1:5, S1vsS2 1:10, S1vsS3 1:50).

## Pseudocode SILAC ratio calculation

### Clean DIA-NN report

1. Filter report.tsv for Precursor.Charge > 1 and remove contaminants

### Create a filter data frame for report.tsv

1. Create an additional column (e.g. “Stripped.Sequence.Charge”) that contains the Precursor.Id without the SILAC label information
2. Filter SILAC data on the reference Channel (H) passing

i. Channel.Q.Value < 0.03
ii. Global.PG.Q.Value < 0.01
iii. Ms1.Translated > 0 and valid value
iv. Precursor.Translated > 0 and valid value Select relevant columns (Run, Stripped.Sequence.Charge, Protein.Group) and save them in a temporary file (“filterSet”) For LFQ data: Filter for Lib.PG.Q.Value < 0.01 and Lib.Q.Value < 0.01
3. Extract information which channels - beyond the reference channel - per Stripped.Sequence.Charge pass the filters described in 2)

a. Inner join filterSet with data (Run, Protein.Group, Precursor.Id, Stripped.Sequence.Charge, Channel.Q.Value & Global.PG.Q.Value, Ms1.Translated, Precursor.Translated)
b. Filter again as in 2), but over all channels
c. Group by Run, Stripped.Sequence.Charge, Protein.Group, Channel
d. Save information which Channels that pass the filters (“Channels.Passed.Standard.Filter”)
e. Overwrite temp file “filterSet”
4. Flag on protein level: How many precursors are used per protein group and in which Channels have the precursors passed the filter

a. Count the number of precursors per protein group and Run in filterSet and save information (includes rescued precursors where only the H channel passed, important for requantify)
b. Count the number of precursors per protein group and Run in filterSet where all channels passed and save information
5. Return this data frame; it should contain the following information on Protein.Group and Precursor level:

i. Run
ii. Protein.Group
iii. Stripped.Sequence.Charge
iv. For each precursor, which channels passed the standard filter
v. How many precursors per protein in total (includes requantify ratios - only reference channel passed filters)
vi. How many precursors per protein where all channels passed filters

### Calculate Protein.Group SILAC ratios and global intensities

1. Create an additional column (e.g. “Stripped.Sequence.Charge”) that contains the Precursor.Id without the SILAC label information (if not already done in before, see above)
2. Create another column “Channel” that contains the channel information from Precursor.Id
3. Select relevant columns (Run, Protein.Group, Channel, Stripped.Sequence.Charge, Ms1.Translated, Precursor.Translated)
4. Create long table where values from Ms1.Translated and Precursor.Translated are in one column
5. Filter for intensities > 0 and valid
6. Create wide table so all intensities are in separate columns per channel
7. Inner join this with filterSet

a. Filter filterSet prior for required number of precursors per protein to pass (either with or without requantify ratios)
b. Only keep precursors that passed the corresponding filter set
c. Only keep precursors where Ms1.Translated and Precursor.Translated are not NA and > 0
8. Calculate Protein.Group L/H ratios by

a. Calculate L/H ratio for every precursor
b. log10-transform those ratios
c. Take the median log10 ratios per protein group and run
9. Calculate L protein abundance

a. Calculate global log10 reference channel abundance

i. Sum the Ms1.Translated and Precursor.Translated intensities of all precursors of a protein for each sample
ii. log10-transform the summed intensities
iii. Take the median of log10 summed intensities over all samples
b. Sum (logged) protein L/H ratios and (logged) global H intensities

### Calculate across-sample ratios

1. Make wide table by samples and Protein.Group intensities calculated in 9) (Samples as columns, L SILAC abundances as values)
2. Calculate across-sample ratios: Sample1 - Sample2, …

a. For LFQ: log10(LFQ Sample 1) – log10(LFQ Sample2)
3. Make sure to not compare a protein where both values only exist thanks to rescued (“requantify”) ratios

## Notes

### Competing Interest Statement

The authors have declared no competing interest.

